# Contrasting genomic evolution between domesticated and wild *Kluyveromyces lactis* yeast populations

**DOI:** 10.1101/2022.09.02.506316

**Authors:** Anne Friedrich, Jean-Sébastien Gounot, Andreas Tsouris, Claudine Bleykasten, Kelle Freel, Claudia Caradec, Joseph Schacherer

## Abstract

The process of domestication has variable consequences on genome evolution leading to different phenotypic signatures. Access to the complete genome sequences of a large number of individuals makes it possible to explore the different facets of this domestication process. Here, we sought to explore the genome evolution of the *Kluyveromyces lactis* yeast species, a well-known species for its involvement in dairy processes but also present in natural environments. Using a combination of short and long-read sequencing strategies, we investigated the genomic variability of 41 *Kluyveromyces lactis* isolates and found that the overall genetic diversity of this species is very high (π = 2.9 x 10^-2^) compared to other species such as *Saccharomyces cerevisiae* (π = 3 x 10^-3^). However, the domesticated dairy population shows a reduced level of diversity (*π* = 7 x 10^-4^), probably due to a domestication bottleneck. In addition, this entire population is characterized by the introgression of the *LAC4* and *LAC12* genes, responsible for lactose fermentation and coming from the closely related species, *Kluyveromyces marxianus*, as previously described. Our results also highlighted that the *LAC4/LAC12* gene cluster was acquired through multiple and independent introgression events. Finally, we also identified several genes that could play a role in adaptation to dairy environments through copy number variation. These genes are involved in sugar consumption, flocculation and drug resistance, and may play a role in dairy processes. Overall, our study illustrates contrasting genomic evolution and sheds new light on the impact of domestication processes on it.

## Introduction

Domestication is a human-related process that shaped the genome of many species. Indeed, by selecting organisms for desirable traits for millennia, humans have acted unconsciously on the evolution of these genomes. Domesticated species therefore represent valuable models for the study of adaptive divergence. While domestication of plants and animals has always been carried out on purpose, the selection for micro-organisms, was first conducted unintentionally, before being better controlled. Indeed, there is evidence for fermented foods and beverages since Neolithic, based first on naturally existing microflora, that predates the use of backslopping technics, far before the discovery of microbes in the late 17^th^ century. In this context, multicellular fungi and yeasts have played a predominant role (Dupont et al., 2017) and access to whole genome sequencing data of large number of individuals having undergone these processes allowed to gain insight into the genomic adaptation footprints at the species level.

Particular interest has been taken in fungal species such as *Aspergillus oryzae*, used for rice and soy fermentation (Gibbons et al., 2012) as well as *Penicillium* fungi such as *P. roqueforti* and *P. camemberti*, both used for the maturation of cheese (Cheeseman et al., 2014; Ropars et al., 2020). Genomic exploration of the latter revealed that their adaptation to cheese environment was associated with recent horizontal transfers of large genomic regions carrying crucial metabolic genes. However, distinct domestication events can co-exist in the same species, generally leading to divergent phenotypic traits (Ropars et al., 2015, 2020). Much of the attention has also been focused on the yeast *Saccharomyces cerevisiae*, a key driver of several industrial fermentation processes. Indeed, the exploration of thousands of complete genomes over the past decade provide evidence for various independent and lineage-specific domestication events within this species, leading to different evolutionary trajectories (Duan et al., 2018; Gallone et al., 2016; Peter et al., 2018; Strope et al., 2015). The beer and bakery populations have been shown to be polyphyletic and diverse at the nucleotide and ploidy level, for example (Bigey et al., 2021; Gallone et al., 2016; Peter, De Chiara, et al., 2018; Saada et al., 2022). In contrast, the wine and sake populations are monophyletic and much less genetically diverse. Genomic processes detected as conferring the desired properties of domesticated strains are diverse and include copy number variant, horizontal gene transfer, large structural variants as well as single nucleotide polymorphisms (Dunn et al., 2012; Gonçalves et al., 2016; Legras et al., 2018; Peltier et al., 2019).

In fact, *Saccharomyces cerevisiae* is no exception and many yeast species have undergone domestication processes (Almeida et al., 2014; Banjara et al., 2015; Hranilovic et al., 2018; Varela et al., 2019). In this context, *Kluyveromyces lactis* represents an interesting species. While it is an attractive model for biotechnological procedures such as production of pharmaceuticals (Spohner et al., 2016), yield of heterologous proteins (Van Ooyen et al., 2006) or bioethanol production (Dickson & Riley, 1989), this aerobic yeast is especially well-known for its ability to ferment lactose. *Kluyveromyces lactis* represents, together with its sister species, *Kluyveromyces marxianus*, one of the leading yeast contributors of dairy products. In *Kluyveromyces lactis*, the lactose assimilation process relies on a well-known pathway comprising the *LAC4* and *LAC12* genes, which encode respectively a β-galactosidase and a lactose permease, as well as the galactose-lactose regulatory genes (*LAC9* and *GAL80*) and the galactose genes (*GAL1, GAL7* and *GAL10*) (Dickson & Riley, 1989; Naumov et al., 2006). This yeast species can also be isolated from natural environments, mainly from insects and tree exudates. These natural isolates are not able to assimilate lactose and ensure its conversion into lactic acid. Based on these physiological and ecological characteristics, the *Kluyveromyces lactis* species was divided into two varieties: *K. lactis* var. *lactis* and *K. lactis* var. *drosophilarum* (Naumov et al., 2014; Sidenberg & Lachance, 1986). The distinction between the two varieties was supported by sequence evidence, as shown by the analysis of the 5.8S-ITS rDNA of multiple samples, but the variety *Kluyveromyces lactis* var. *drosophilarum* is also very heterogeneous and can be divided into several populations (Naumova et al., 2004). Based on a study of 12 isolates, it was shown that the *LAC4* and *LAC12* genes are found specifically in the var. *lactis* strains and confer the ability to ferment lactose (Naumov et al., 2006). Recently, it has been established that these two genes were acquired by introgression from *Kluyveromyces marxianus* (Varela et al., 2019). A population genomic study based on the analysis of the complete genome of 14 *Kluyveromyces marxianus* isolates revealed that dairy and non-dairy strains differ mainly by polymorphism within the *LAC12* gene but also by variation in ploidy level, suggesting that multiple characteristics separate the dairy and non-dairy strains (Ortiz-Merino et al., 2018).

To date, the evolutionary history of the *Kluyveromyces lactis* species remains unclear, and the comparative genomic analysis of the wild and dairy strains was based on a limited number of isolates and only focused on short genomic regions, preventing a complete view of genome evolution and the impact of domestication within this species. We therefore sought to study the evolutionary history of this species based on the whole genome sequence analysis of 42 *Kluyveromyces lactis* strains, originating from both dairy and non-dairy environments. As an interesting alternative model in yeast genetics, *Kluyveromyces lactis* was among the first eukaryotic organisms to have its complete genome sequenced (Dujon et al., 2004). The CBS 2359 reference strain, which belongs to the variety *Kluyveromyces lactis* var. *lactis*, was chosen as reference and the availability of this high quality 10.6 Mb sequence and its associated annotations allows the establishment of a population genomic analysis. Exploration of the global pattern of polymorphisms allowed to draw a precise view of the phylogenetic relationships between strains, revealing very separated populations in which individuals are very closely related. Our analyses confirm that the entire dairy population is characterized by an introgression of the *LAC4* and *LAC12* genes from *Kluyveromyces marxianus*. However, careful determination and examination of the structure of the introgressed regions revealed that this gene cluster was acquired through multiple and independent introgression events after the divergence of the dairy and wild population. In addition, using copy number variants, we identified several genes whose presence/absence pattern could indicate a role in adaptation to dairy environments. Overall, our study offers new insights into the evolutionary history of *Kluyveromyces lactis* and the impact of domestication processes on it.

## Results

### *Kluyveromyces lactis* is characterized by a high genetic diversity and a structured population

For this study, we gathered a collection of 42 isolates from around the world and coming from diverse ecological niches (Table S1). Within our collection, 19 strains have been isolated from dairy environments (*e.g*. cheese, buttermilk or cream) and come mainly from Europe. Natural isolates have been mostly isolated from insects and trees in Asia and North-America. We subjected the almost entire collection (41 strains out of the 42, the last one corresponding to the type strain previously sequenced) to short-read whole-genome sequencing, with a mean coverage of 115-fold per sample. The reads associated with each of the 41 samples were mapped to the *Kluyveromyces lactis* reference genome. A total of 1,767,970 reference-based polymorphic positions were detected in the population, 1,594,125 being related to SNPs (Single Nucleotide Polymorphims) and 173,845 to small indels.

The SNP dataset was used to evaluate the overall genetic diversity within the species. The average pairwise difference between strains π reaches 2.8 x 10^-2^, which is almost 10-fold higher compared to *S. cerevisiae* (3 x 10^-3^) (Peter et al., 2018). This population-level genetic divergence is, to our knowledge, the highest reported to date within a yeast species. While several species such as *Saccharomyces uvarum* and *Lachancea kluyveri* showed greater diversity compared to *S. cerevisiae* (1.2 x 10^-2^ and 1.7 x 10^-2^, respectively), none were ever mentioned as exceeding 2 x 10^-2^ (Almeida et al., 2014; Friedrich et al., 2015). Interestingly, the diversity within the closely related species, *Kluyveromyces marxianus*, in which dairy and wild strains also coexist, is much lower and has been estimated at π = 1.2 x 10^-2^ based on the sequencing data generated for a collection of *Kluyveromyces marxianus* genomes (Ortiz-Merino et al., 2018).

This SNP dataset was used for a neighbor-net (SplitsTree) analysis that clusters the strains into five highly separated populations (Figure 1A), within which individuals are closely related (Table S2). This topology was confirmed by the inference of the phylogenetic relationships between the strains, through the construction of a maximum-likelihood tree, as well as a neighbor-joining tree (Figure S1). The largest population corresponds to the *Kluyveromyces lactis* var. *lactis* part of the tree and includes 21 strains among which the reference strain and all those associated with dairy products. Excluding UCD 70-4, which was isolated from a winery in South Africa and is significantly more divergent, the average pairwise divergence observed between the strains in this group is 0.06% (Table S3). The very low genetic divergence in this population is very close to what can be observed, for example, in the sake population of *Saccharomyces cerevisiae*, with an average divergence of 0.08%, which may be a distinctive sign of domestication (Peter et al., 2018). The *Kluyveromyces lactis* var. *drosophilarum* strains are distributed in four distantly related populations whose segregation correlates mostly with the geographical origins of the strains (Figure 1A, Figure S2). We named these clusters based on these origins, leading to an Asian cluster and three distinct North-American clusters (NA1, NA2 and NA3). The intra-group mean genetic divergence ranges from 0.26 to 0.78 for the NA2 and NA3 populations, respectively, which is much higher compared to the dairy population. The population structure is in complete accordance with the distribution of the strains on the tree and each group is represented as a clear population for which no admixture has been highlighted (Figure 1, Figure S3). This suggests the absence of interlineage outcrossing events, which could be attributed to geographical separation of the isolates. These results confirmed the great heterogeneity within *Kluyveromyces lactis* var. *drosophilarum* and the existence of clear populations, already suggested by the analysis of small genomic regions (Naumova et al., 2004).

**Figure 1:**
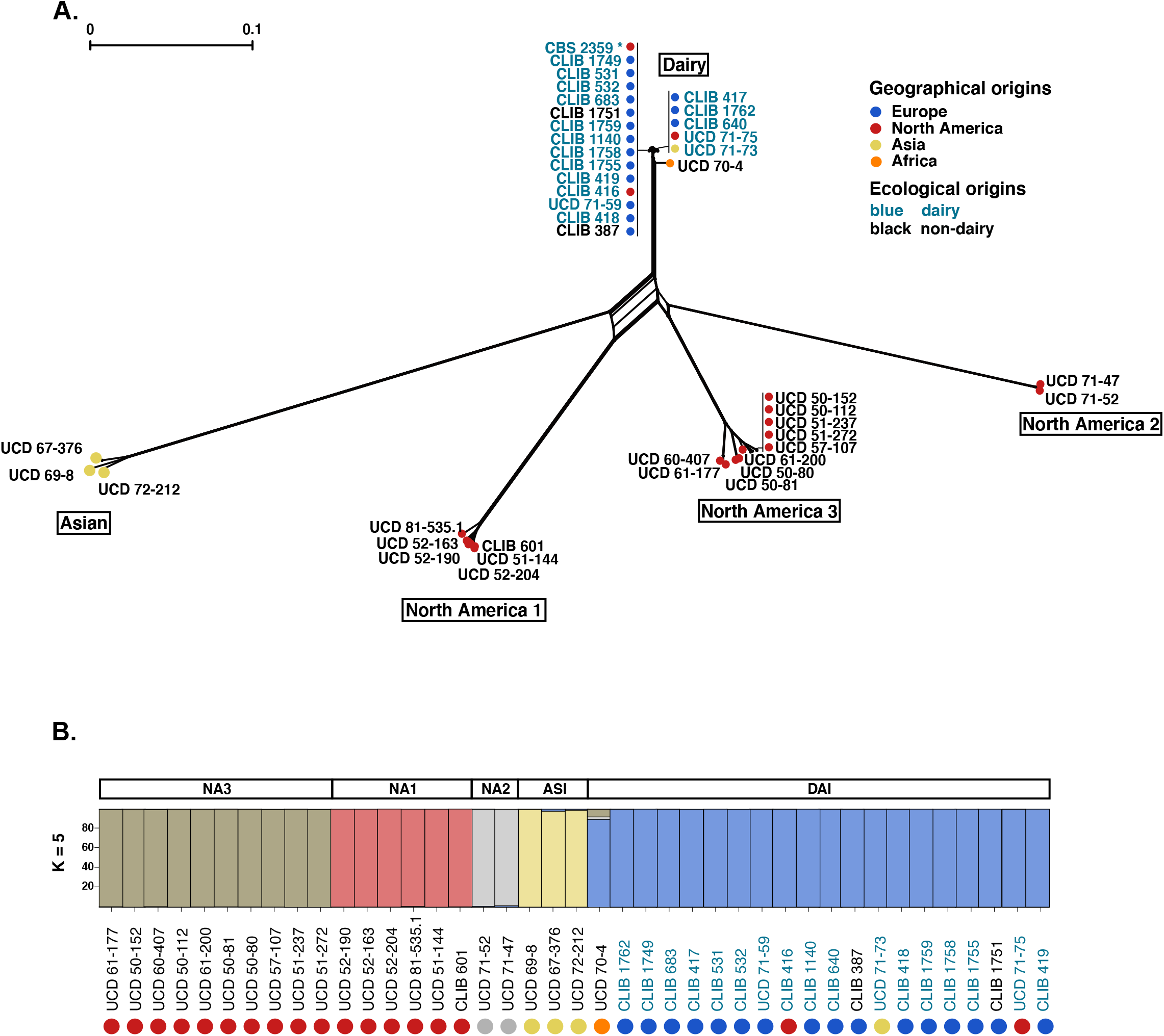
Population structure of the 42 *Kluyveromyces lactis* studied strains. **A.** Neighbor-net analysis based on 1,594,125 SNPs identified in the surveyed strains. Branch lengths are proportional to the number of sites that discriminate each pair of strains and cross-linking indicates the likely occurrence of recombination. **B.** Structure of the population, with K=5 populations. Each strain is represented by a vertical bar, which is divided into segments that represent the strain’s estimated ancestry proportion in each of the 5 populations. The circle colors denote the geographical origins of the strains.

Interestingly, the genetic divergence between the clusters is very high, exceeding 8.5 SNPs every 100 bp between the two most divergent populations, namely the Asian and North America 2 groups (Table S4). Such genetic diversity between the populations along with the absence of reticulation within the tree (Figure 1A) raised the question of reproductive isolation within this species. In a previous study, isolates from different populations were crossed and the spore viability was estimated (Naumov & Naumova, 2002). While the intra-population crosses led to fertile hybrids, the inter-cluster crosses led to very low viability (< 45 %). The recent release of the complete genome of the type strain of *Kluyveromyces lactis* var. *drosophilarum* (CBS 2105 = CLIB 601), which belongs to the NA1 cluster, showed large chromosomal rearrangements compared to the type strain of *K. lactis* var. *lactis* (CBS 2359 = CLIB 210), the reference isolate of the dairy population (Varela et al., 2019). These types of evolutionary events could lead to reproductive isolation, as could the strong genetic divergence associated to the allopatric distribution of taxa that could lead to incompatible alleles. Taken together, these observations raise the definition of *Kluyveromyces lactis* as a species in itself and likely point to an early stage of a speciation process.

### Ploidy level and chromosome number variation

Systematic analysis by flow cytometry confirmed that the *Kluyveromyces lactis* species is primarily a haplontic species because all strains have a haploid profile associated with it, with the exception of UCD 57-107 for which a diploid profile was obtained (Table S1). UCD 57-107 showed a very low proportion of heterozygous SNPs (<1%), comparable to what was observed for haploid strains, and was therefore considered as homozygous. The read coverage profiles along the reference genome revealed that our collection was devoid of aneuploidy. Based on 10-kb sliding windows, a total of 14 segmental duplications were detected in seven isolates (from 1 to 7 segmental duplications per isolate) (Table S1), all chromosomes being affected but chromosome 2. Chromosome 4 was the most impacted, with 4 strains carrying a segmental duplication on the right arm of this chromosome. The lack of variability in the level of ploidy in *Kluyveromyces lactis*, as well as the absence of variability of the number of chromosomes contrast with what has recently been observed in other yeast species. In the case of its sister species, *Kluyveromyces marxianus*, the variation of ploidy level has been shown to separate dairy and non-dairy strains, the dairy strains being all diploid or triploid, whereas non-dairy strains being haploid (Ortiz-Merino et al., 2018). Moreover, higher ploidy level and aneuploidies were also recently described as enriched in some domesticated clades of *S. cerevisiae* (Peter, De Chiara, et al., 2018). Altogether, our results suggest that these mechanisms are not driving evolutionary processes related to dairy environment in *Kluyveromyces lactis*.

### Gene content, copy number variation and domestication

The *LAC4* and *LAC12* genes, that encode respectively for β-galactosidase and lactose permease, have been identified for more than a decade as controlling the fermentation of lactose in *Kluyveromyces lactis* var. *lactis* as these two genes were detected as absent in strains unable to ferment lactose (Naumov et al., 2006). Here, the presence of these genes was investigated at the whole collection level, and as expected, they were confirmed to be absent in all the strains that did not belong to the dairy cluster (Figure 2A). Within the dairy cluster, a single strain was detected as not carrying these genes, the UCD 70-4 isolate. This latter was isolated in a winemaking equipment and was already described as closely related to dairy strains but without dairy capabilities (Naumov et al., 2006). This characteristic could be attributed either to a loss of these genes or to the absence of introgression events. The outlier position in the dairy branch of the tree suggests that this isolate might actually be a close relative to the dairy ancestor that underwent the introgression event.

**Figure 2:**
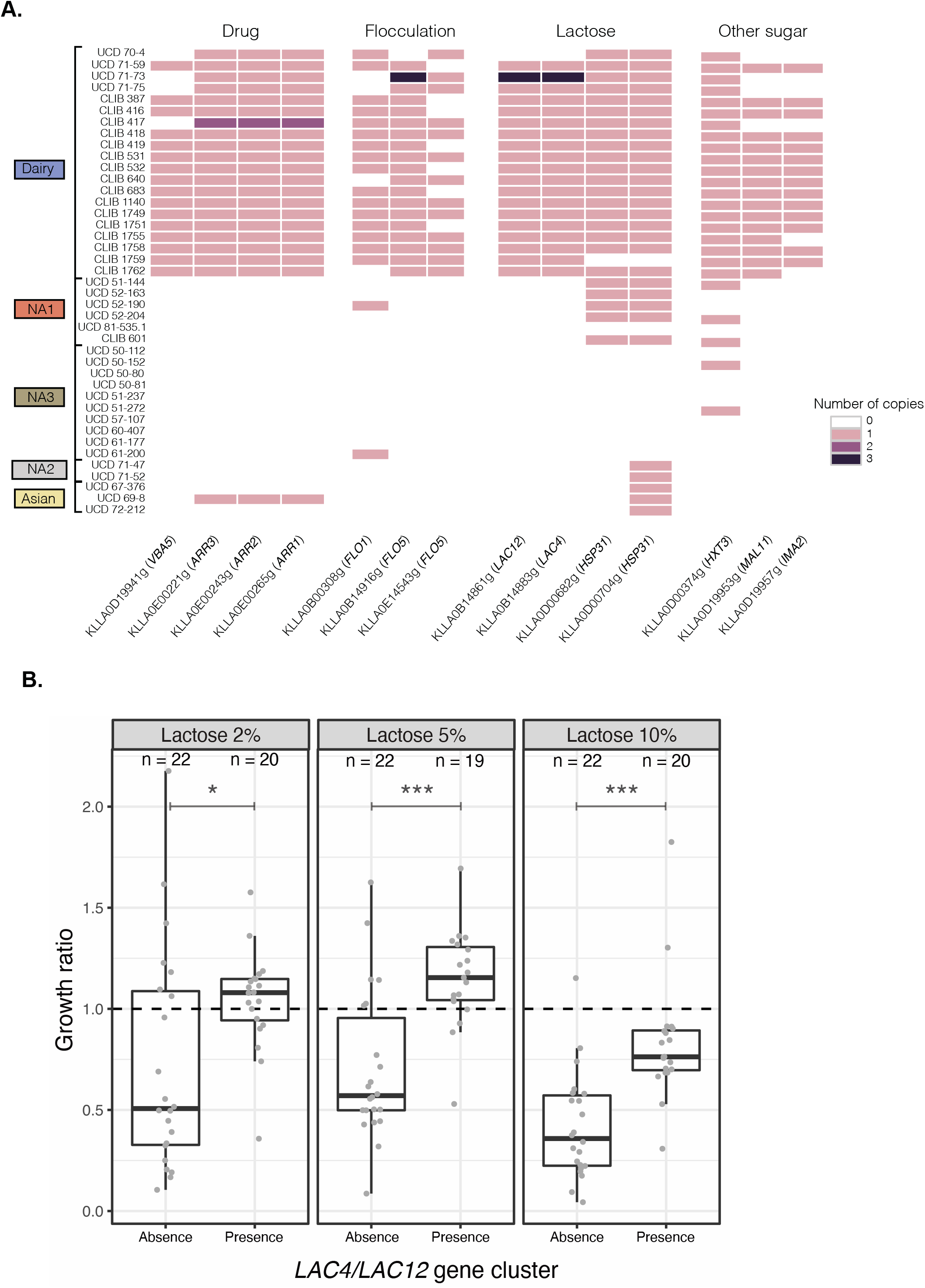
Diversity in terms of gene copy number within the population. **A.** Number of copies of a subset of genes having differential profiles between the dairy and non-dairy isolates. Strains are organized by population. **B.** Distribution of the growth ratio, *i.e*. the max OD in the tested conditions normalized by the max OD obtained in glucose 2% condition, of the *Kluyveromyces lactis* strains with increasing concentration of lactose, according to the presence or the absence of the *LAC4/LAC12* gene cluster. Wilcoxon–Mann–Whitney test was applied to assess the significance of the growth ratio difference between the two groups of strains. The level of significance is indicated as follows: ns: not significant, *P ≤ 0.05, **P ≤ 0.01, ***P ≤ 0.001.

To test the ability of our strain collection to assimilate lactose, the growth of the 42 natural strains was recorded in complete medium alone and with increasing concentration of lactose (2%, 5% and 10%), using a microcultivation approach. These results clearly demonstrate better growth capacities under lactose conditions for the strains having the *LAC4* and *LAC12* genes, this difference being more significant with the increase in the lactose concentration (Figure 2B). This growth advantage is most likely related to the presence of this gene cluster, however we cannot exclude that some other genes contribute to this phenotype.

While the main driver of dairy/non-dairy properties is obviously captured by the differential presence of this gene cluster, the availability of sequencing data for a large *Kluyveromyces lactis* collection have paved the way for a more systematic view of the variants that discriminate populations at the genomic level. To this end, gene copy number variants (CNVs) were first investigated based on read mapping to the *Kluyveromyces lactis* var. *lactis* reference with Control-FREEC (Boeva et al., 2012). Within the whole population, we detected 1,406 gene gains and 2,027 gene losses related to 937 unique genes annotated on the reference genome (Figure S4A). However, the distribution of CNVs is highly impacted by segmental duplications, in particular for the UCD 50-81 and UCD 50-80 strains, for which 270 and 146 specific gene gains are respectively reported (Figure S4B). To identify genes with differential profiles between dairy and non-dairy populations, we applied a two-sample Kolmogorov-Smirnov test and selected genes with a p-value lower than 0.05. This leads to the identification of 87 genes, most of which share a common profile, *i.e*. predominantly present in the dairy population and absent in the others (Table S5). Among this set of genes, *LAC4, LAC12* and *FLO5*, part of the regions introgressed from *Kluyveromyces marxianus*, are the genes that are found systematically in the dairy strains only. A single strain, UCD 71-73, showed a higher copy number of these genes, suggesting that duplication of these genes is not a driving factor for dairy environments adaptation. In addition, some interesting genes are also present in this set of 87 genes. For example, two genes, KLLA0D00682g and KLLA0D00704g, are annotated as similar to the *HSP31* gene of *S. cerevisiae*, known to be involved in oxidative stress resistance and which methylglyoxalase activity converts methylglyoxal to D-lactate (Tsai et al., 2015). Certain genes involved in the metabolism of other sugars and in flocculation such as the maltose transporter *MAL11*, the isomaltase *IMA2* and the hexose transmembrane transporter *HXT3* were also detected in most of the dairy strains although they were not found in the majority of wild strains. These genes have already been found in bread and cheese *S. cerevisiae* strains and also play an important role during dairy processes (Legras et al., 2018). Finally, genes involved in drug resistance, including the plasma membrane protein *VBA5* and the *ARR* genes (*ARR1-3*) linked to the arsenate resistance, showed similar profiles. These genes have already been identified as playing a major role in arsenate resistance in *S. cerevisiae* and could therefore play a role during industrial processes (Bobrowicz et al., 1997; Peter, De Chiara, et al., 2018).

### Multiple introgression events of the *LAC* cluster in the dairy population

While the differential presence of the *LAC4/LAC12* gene cluster has been identified for over a decade as controlling the fermentation of lactose in *Kluyveromyces lactis* var. *lactis* (Naumov et al., 2006), it was only recently shown that this gene cluster was acquired by introgression from the sister species *Kluyveromyces marxianus* (Varela et al., 2019). This event also leads to the integration of a third gene, namely *FLO5*. To better understand the structure and the evolution of this introgressed region, we sought to completely assemble the genomes of all strains of the dairy population, based on our Illumina sequencing data. As expected, the presence of this three-gene cluster was detected in all Illumina assemblies except UCD70-4, that stands at the root of the dairy part of the tree and was isolated in a South African winery. Two scenarios can be considered here: either the acquisition of the introgressed region occurred after the divergence between the ancestor of the UCD 70-4 and the other dairy strains which would imply that the divergence between *Kluyveromyces* var. *drosophilarum* and var. *dairy* populations preceded the introgression event, or this strain lost the introgressed region during its evolutionary history.

Surprisingly, the presence of an additional gene coming from *Kluyveromyces marxianus*, *CEL2*, was detected next to the introgressed regions for 5 isolates, namely CLIB 417, CLIB 640, CLIB 1762, UCD 71-73 and UCD 71-75. The *CEL2* gene is the first neighbor of *LAC12* on the *Kluyveromyces marxianus* genome from which the introgressed region originates (Figure 3). This observation raised the question of the evolutionary history of introgressed regions and whether a larger region encompassing *CEL2* was first introgressed into the common ancestor of dairy strains and then lost in some isolates, or whether multiple introgressions have occurred within the population. The very low sequence divergence of this region in the dairy strains does not allow us to conclude on the basis of this criterion. In this context, we sought to explore the genomic context of these regions. The high level of fragmentation of Illumina assemblies prevents us from having a clear view of the genomic structure within these isolates. This motivated long-read sequencing of six isolates from the dairy population: two strains harboring the *CEL2* gene (CLIB 640 and UCD 71-75), three strains lacking this gene (CLIB 1751, CLIB 1759 and CLIB 419), as well as the non-dairy strain UCD 70-4 (Table S1). Chromosome-level assemblies were generated for all of them and made it possible to highlight two different genomic structures for this introgressed region (Figure 3). Indeed, the three-genes cluster shared all the same chromosomal location as in the reference strain, *i.e*. at the end of the right-arm of chromosome 2 (version v1), while the four-genes clusters were detected at the end of the right-arm of chromosome 3 for both strains (version v2) (Figure 3). Comparative analysis of these chromosomal structures suggested that several independent introgression events could have occurred from *Kluyveromyces marxianus* into the *K. lactis* dairy genomes. Indeed, the end of the right arm of chromosomes 2 are entirely syntenic between UCD 70-4 and the *Kluyveromyces lactis* var. *lactis* v2 background. This region is also mostly syntenic with *Kluyveromyces lactis* var. *drosophilarum* (Varela et al., 2019), the difference being based on the presence of 10 additional genes in the very subtelomeric regions that are known to be highly variable in yeast genomes. It should be noted that the synteny between the *FET5* and *KLLA0B14839* genes is conserved in these three cases. Consequently, in case of a single introgression event of the *LAC* cluster within the dairy group, the one observed in *Kluyveromyces lactis* var. *lactis* v2 background should have been the ancestral event and would have preceded the accumulation of chromosomal rearrangements that would have led to the *Kluyveromyces lactis* var. *lactis* v1 chromosomal structure. Nevertheless, the complete synteny observed at the end of the right arm of chromosome 3 between UCD 70-4 and *Kluyveromyces lactis* var. *lactis* v1 background tends not to support this scenario and allowed us to propose that at least two independent introgression events occurred within the *K. lactis* var. *lactis* population.

**Figure 3:**
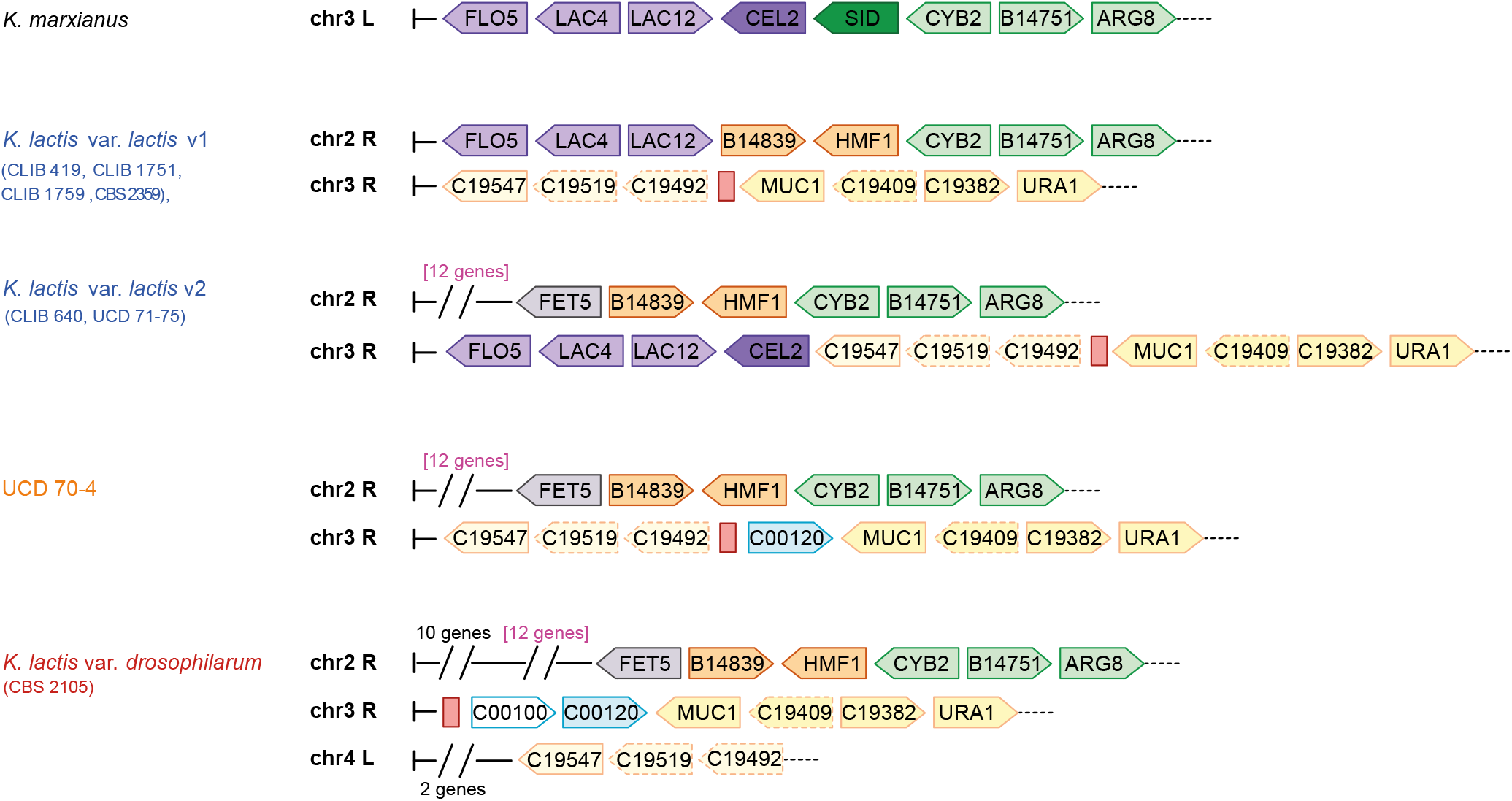
Chromosomal structure of the introgressed *LAC4/LAC12* gene cluster in several *Kluyveromyces lactis* and *K. marxianus* backgrounds. On this schematic representation of the chromosomal regions related to the introgression events that occurred from *Kluyveromyces marxianus* into the genome of the *K. lactis* dairy strains, *K. lactis* var. *lactis* v1 referred to the dairy reference strain, *K. lactis* var. *lactis* v2 referred to the newly detected cluster in some dairy strains. The name of the strains is colored according to their origins: blue for European dairy, orange for African non-dairy and red for North-American non-dairy. L and R referred respectively to the left and right extremities of the chromosomes. Vertical bars (on the left) represent the end of the chromosomes, while the black dotted lines (on the right) indicate that the chromosomes continue. Genes in syntenic regions share the same color and dotted boxes are associated to pseudogenes. Red boxes indicate the presence of LTR elements.

Finally, the synteny observed at the right-end of chromosomes 2 as well as at the right-end of chromosomes 3 between UCD 70-4 and *K. lactis* var. *lactis* v2 support the fact that the divergence of the so-called dairy and the other clades predates the acquisition of dairy properties. In addition, UCD 70-4 is at the forefront of this acquisition and it can therefore be defined as the closest relative to the ancestral strains into which the introgressions occurred.

### Transposable element content is clade specific

It has recently been shown that the evolution of the transposable element (TE) content follows the population structure in *S. cerevisiae* and is deeply impacted by clade-specific events such as introgressions (Bleykasten-Grosshans et al., 2021). The genome-wide content of transposable element was investigated through short-read mapping on finely selected TE queries originating from both *Kluyveromyces lactis* and *K. marxianus* species. The TE landscape clearly shows that the *Kluyveromyces lactis* populations display specific TE repertoires, which are mainly defined by population specific TE (Figure S5). When a TE is present in a given isolate, it is found in most of the isolates of the related population and absent in the other populations. For example, a TE family similar to Tkl-3 query is specific to the Asian groups, whereas TE families similar to Tkl-1, Rkl-3 or Rkl-4 are specific to dairy isolates. This exclusive distribution of population specific TEs that are unrelated contrasts with the population-specific landscape that has been observed in *S. cerevisiae*, which relies on both population-specific TEs from the same TE family (*i.e*. population specific Ty1 sequence variants) and population specific combinations of TEs shared by multiple populations. Interestingly, none of the *Kluyveromyces marxianus* TE were detected in the whole population. In addition, the dairy population carries the lowest TE content. In most cases, only one copy per genome is present, and in many strains, the Tkl-1 and Tkl-2 queries highlight only partial coverage, suggesting either that the corresponding TE are truncated or that they present segments of sequence too divergent to allow the mapping of reads. There is a striking exception represented by the CLIB418 isolate which contains about a hundred of Tkl-1 related copies. These copies are clearly the result of a transposition burst, as supported by the uniform shape of the coverage profile indicating that they are very similar.Mitochondrial genome diversity in *K. lactis*

The mitochondrial genome of *Kluyveromyces lactis* is 40.3 kb long and contains the same set of eight protein-coding genes as *S. cerevisiae*. These genes encode three subunits of the ATP synthase complex (*ATP6, ATP8, ATP9*), the apocytochrome b (*CYTB*), three subunits of the cytochrome c oxidase complex (*COX1, COX2, COX3*) and the ribosomal protein (*VAR1*). We sought to explore mitochondrial genome diversity within our collection through genome assemblies, but their high level of fragmentation prevents comparison at the whole mitochondrial genome level. Complete gene coding sequences could only be recovered for 4 genes (*ATP6, COX2, COX3* and *CYTB*) across the 41 isolates. Interestingly, the *ATP6* and *COX2* genes were recently identified as the most informative to describe *S. cerevisiae* population structure and phylogeny (De Chiara et al., 2020). So, we confidently thought that limiting our analysis to these 4 genes would still help to better understand mitochondrial genome evolution.

The genetic diversity related to these four genes was lower compared to the nuclear coding genes, for which a mean π value of 0.023 was observed. However, this mitochondrial genetic diversity varies a lot according to the considered gene (Table S6): *COX2* and *ATP6* showed very low genetic diversity, with *π* values of 0.0011 and 0.0036, respectively, while *COX3* and *CYTB* had much higher genetic diversity, with π values of 0.0160 and 0.0165. These results are in contrast with what was observed in *S. cerevisiae* for which *COX2* and *ATP6* were among the highly divergent genes (with π values of 0.0166 and 0.0108, respectively) and *COX3* and *CYTB* were among the lowly divergent genes (with π values of 0.0072 and 0.0048, respectively). These four gene sequences were concatenated and used to construct a neighbor-joining tree (Figure S6). Interestingly, the phylogeny recapitulates quite efficiently most of the populations observed at the nuclear level, with the exception of isolates from the two North-America groups, which were completely mixed, revealing some mito-nuclear discordance (Figure S6). This contrasts with what was recently observed for *S. cerevisiae* (De Chiara et al., 2020), for which a very discordant evolution of the mitochondrial and nuclear genomes was observed, due to outbreeding and recombination events between the parental mitochondrial genome. These results suggest the absence of natural crosses between isolates from different populations, with the exception of North American strains which may not have been geographically isolated.

## Conclusion

Overall, our study represents the first population genomic survey focusing on a large population of *Kluyveromyces lactis* isolates. This species is of interest as it is partially domesticated, which leads to differential genome evolution across the various identified populations. By sequencing the whole genome of 41 *K. lactis* isolates (*Kluyveromyces lactis* var *lactis* and var *drosophilarum*), we found the highest intraspecific genetic divergence ever reported for a yeast species. Compared to other yeast species explored to date, *K. lactis* presents a nucleotide diversity (estimated at 2.8 x 10^-2^), which is on average 2-fold higher (*e.g*. for *S. cerevisiae, S. uvarum, L. kluyveri, π* is approximately ~ 4 x 10^-3^, ~1.2 x 10^-2^ and ~1.7 x 10^-3^, respectively) (Schacherer et al., 2009; Almeida et al., 2014; Friedrich et al., 2015). Based on the SNP diversity, the isolates grouped into five distinct clusters with the dairy group (*Kluyveromyces lactis* var *lactis*) being well separated from the wild ones (*Kluyveromyces lactis* var *drosophilarum*). However, the four wild populations are also very distant as the divergence between these clusters is above 3 % and can exceed 8 % between the most divergent ones, namely the Asian and North America 2 groups. These observations raise the definition of the *Kluyveromyces lactis* species and probably highlights an early stage of a speciation process. This hypothesis is also supported by the presence of reproductive isolation between all these populations, as previously described. While the intra-population crosses led to fertile hybrids, the inter-cluster crosses led to very low viability (< 45 %) (Naumov & Naumova, 2002).

Our study also provided a better understanding of the domestication process of the dairy population of *K. lactis (Kluyveromyces lactis* var *lactis*). Recently, there has been a particular interest in exploring the domestication processes of different species found in milk and dairy products such as *Kluyveromyces marxianus* (Ortiz-Merino et al., 2018) as well as used for the maturation of cheese such as *P. roqueforti* and *P. camemberti* (Cheeseman et al., 2014; Ropars et al., 2020). Even if certain characteristics are common to these different processes of domestication, they all have their specificity. Here, we found that the nucleotide variability within the *Kluyveromyces lactis* dairy population is very limited. Indeed, the maximum pairwise diversity ranges from 0.23 % to 0.79 % for the wild clusters, whereas it does not exceed 0.06 % between the strains presenting dairy capacities. This reduced genetic diversity within the dairy cluster seems to be a clear feature of the domestication process in *K. lactis*.

As already shown for species isolated from dairy products or used in cheese maturation, horizontal gene transfers or introgressions are key players of domestication events (Cheeseman et al., 2014; Ropars et al., 2020; Varela et al., 2019). Although the impact of the introgression of the *LAC4/LAC12* genes cluster from *Kluyveromyces marxianus* was already assessed, our study also highlighted some interesting evolutionary features regarding this aspect in the *K. lactis* dairy population. We first showed that the divergence between the *Kluyveromyces lactis* var*. lactis* and the *K. lactis* var*. drosophilarum* isolates occurred before the acquisition of the dairy abilities, i.e. the introgression of the *LAC4/LAC12* genes cluster. Our results also suggest that the UCD70-4 strains, which is part of the dairy cluster without dairy capabilities, should be close to the ancestral strains in which introgressions occurred. Additionally, the genomic resolution we obtained using long-read sequencing and *de novo* assemblies allowed us to suggest that several independent introgression events occurred within this species.

Finally, it should also be noted that in *Kluyveromyces lactis*, no real variability in terms of ploidy was observed, and aneuploidies and CNVs are not predominant. This observation contrasts with other yeast species that have undergone domestication processes, in particular with *Kluyveromyces marxianus*, for which ploidy variation separates the dairy and non-dairy strains (Ortiz-Merino et al., 2018), but also *S. cerevisiae* for which some domesticated populations, such as the beer isolates, have been described as having a higher ploidy level (Peter et al., 2018; Saada et al., 2022).

Overall, our study sheds new light on the domestication of the dairy population of *Kluyveromyces lactis*. In addition, it also clearly shows once again that even some common characteristics might be shared, each domestication process is specific and unique to each domesticated population.

## Materials and methods

### Studied strains

A collection of 42 *Kluyveromyces lactis* strains was compiled for this study (Table S1) with the aim to maximize the ecological and geographical origins of the strains. The samples were mostly purchased from the Phaff and the CIRM collections. Our final dataset contains 17 strains directly related to dairy processes, that originates mostly from Europe. The wild strains were mostly isolated from insect and tree exudates in North America and Asia.

### Illumina sequencing and polymorphisms detection

To obtain genomic DNA for 41 strains sequenced in the context of this study, isolates were grown overnight at 30°C in 20 mL of YPD medium to early stationary phase before cells were harvested by centrifugation. Total genomic DNA was subsequently extracted using the MasterPure Yeast DNA purification kit (Cat No MPY80200) according to the manufacturer’s instructions. For all genomes, 280-bp insert libraries were produced and sequenced on an Illumina HiSeq 2000 platform. 100-bp paired-end reads were generated.

For each strain, reads were mapped against the reference genome of the *Kluyveromyces lactis* var. *lactis* type strain CBS 2359 (GenBank accession numbers CR382121 to CR382126) using BWA mem v.0.7.15 (Li & Durbin, 2009) with default parameters. Alignments were then successively cleaned using samtools fixmate function v.1.3.1 (Li et al., 2009), GATK v.4.0.11 realignment function (McKenna et al., 2010) and Picard MarkDuplicates function v.1.140 (broadinstitute.github.io/picard). The obtained BAM files were then used to detect single nucleotide polymorphisms and small indels using GATK HaplotypeCaller with a requested ploidy set to one. GVCF files obtained were finally concatenate into one single GVCF using GATK CombineGVCFs and GenotypeGVCFs functions, allowing to produce correct genotype likelihoods and re-genotype the newly merged record.

### Oxford Nanopore sequencing

Yeast cell cultures were grown overnight at 30°C in 20 mL of YPD medium to early stationary phase before cells were harvested by centrifugation. Total genomic DNA was then extracted using the QIAGEN Genomic-tip 100/G according to the manufacturer’s instructions. The extracted DNA was barcoded using the EXP-NBD104 native barcoding kit (Oxford Nanopore) and the concentration of the barcoded DNA was measured with a Qubit^®^ 1.0 fluorometer (Thermo Fisher). The barcoded DNA samples were pooled with an equal concentration for each strain. Using the SQK-LSK109 ligation sequencing kit (Oxford Nanopore), the adapters were ligated on the barcoded DNA. Finally, the sequencing mix was added to the R9.3 flowcell for a 48 hour run. Basecalling was performed with Guppy (v2.3.5) and raw fastq files were treated with porechop (v0.2.3) to remove both adapters and barcodes.

### Tree building, divergence calculation and structure analysis

We inferred phylogenetic relationships among the 41 isolates using the dataset of 1,594,125 SNPs in a maximum likelihood framework with IQ-Ttree2 (Minh et al., 2020), with TVM+F as model. This latter was chosen as best-fit model by BIC through ModelFinder (Kalyaanamoorthy et al., 2017). These polymorphic positions were also used for the neighbor-net analysis which was performed with SplitsTree5 software via the SplitsNetworkViewer.

The isolate relationships, were also assessed through the construction of a distance tree using the BioNJ algorithm (Gascuel, 1997) provided in the SplitsTree4 software (Huson & Bryant, 2006). To that end, a sequence representative of each strain was constructed by inferring SNPs within the reference chromosomes, that were than merged in a single 10.7 Mb sequence. All strains sequences were given as input to SplitsTree.

These sequences were also leveraged to estimate the pairwise divergence between each strain as the ratio between the number of non-equal positions between the two considered strains and the total size of the genome.

Finally, these sequences were also used for the inference estimation of the number of population clusters through STRUCTURE software, version 2.3.4 (Pritchard et al., 2000). We ran the software independently with a number of populations K ranging for 2 to 6 using the admixture model with a burn-in period of 100,000 and 500,000 replicates.

### Calculation of population genetic statistics

In order to get an estimate of the nucleotide diversity at population level, *π*, the average pairwise nucleotide diversity θw, the proportion of segregating sites and Tajima’s D value, which represents the difference between *π* and θw were computed. Multiple alignments of the concatenated chromosomes that were representative of the isolates were submitted to variscan (Vilella et al., 2005) with options runmode set to 12 and usemuts set to 1.

*π* was also estimated for the *Kluyveromyces marxianus* species, based on the Illumina reads generated in the context of (Ortiz-Merino et al., 2018) and the reference sequence of the species (GenBank accession numbers AP014599 to AP014607).

### Determination of copy number variants

The detection of copy number variants (CNVs) related to the *Kluyveromyces lactis var. lactis* reference and affecting each strain was performed by running Control-FREEC (Boeva et al., 2012), version 10.6 on their respective BAM files. The program was used with the following parameters: breakPointThreshold = 0.6, window = 1000, telocentromeric = 600, step = 200, ploidy = 1, minExpectedGC = 0.3 and maxExpectedGC = 0.4. To obtain a count of the number of copies for each genomic feature, Control-FREEC output files were crossed with the reference genome annotations. Features for which at least half of the length was contained in a region whose number of copy deviates from one were considered under CNV. If regions with different coverages overlapped a single feature, the one covering the larger part of the feature was considered.

### Analysis of the copy number variants

To identify genes with variable CNV patterns between the dairy and non-dairy varieties, a two-sample Kolmogorov-Smirnov statistic test was applied to each of the 5,076 protein coding genes annotated in the reference genome. Genes for which distributions differ between both varieties (p-value < 0.05), were further selected for manual inspection.

### Transposable element detection

The strategy used in (Bleykasten-Grosshans et al., 2021) was adapted for this study. Briefly, a set of 23 sequences originating from *Kluyveromyces lactis* and *K. marxianus* was defined as the representative dataset. Among them, nine sequences belong to the Class II Rover elements from the hAT superfamily described in (Sarilar et al., 2015). The other 14 query sequences are similar to Class I LTR elements from the Copia superfamily. The Class I query sequences were named Tkl or Tkm (for Ty *Kluyveromyces lactis* and Ty *Kluyveromyces marxianus*, respectively), according to the nomenclature used in (Neuvéglise et al., 2002). Similarly, the Class II sequences were named Rkl or Rkm (for Rover *Kluyveromyces lactis* and Rover *Kluyveromyces marxianus*, respectively). The Illumina reads of the 41 strains were mapped against this dataset and the coverage profiles were manually inspected. Only eight *Kluyveromyces lactis* queries showed significant coverage, that allowed estimating the number of copies of the considered element in the different strains.

### Whole genome assembly construction

Illumina paired-end reads assemblies were constructed with Abyss (v.2.0.2) (Simpson et al., 2009) with the option ‘-k 64’.

For the long-read assemblies, the Oxford Nanopore fastq files were downsampled with Filtlong (v0.2) (https://github.com/rrwick/Filtlong) to get a 40X coverage per strain (options --min_length 1000 -- mean_q_weight 10). The downsampled datasets were than independently assembled with with SMARTdenovo (Liu et al., 2021), with the options ‘-c 1 -k 16 -J 5000 -e zmo’.

### Strain phenotyping and growth quantification

The 42 *Kluyveromyces lactis* strains were inoculated into flat bottom 96-well microplates (Nunclon, Thermo Fisher) containing 150 μL of YPD (Yeast extract 1% Peptone 2% Dextrose 2%) and incubated overnight at 30 °C. Pre-cultures were washed 5 times in sterile MilliQ water to eliminate residual glucose. After the last washing step, cells were resuspended in sterile MilliQ water and transferred with a Micro-Plate Pin Replicator in a 96-well microplate containing 150 μL of synthetic complete media (yeast nitrogen base with ammonium sulfate 6.7 g/L, SC amino acid mixture 2 g/L) with either glucose (2 %) or lactose (2 %, 5 % or 10 %) as carbon source. Optical density (OD) at 600 nm was measured every ten minutes during 48 h at 30 °C using the microplate reader TECAN Infinite^®^ 200Pro. Before each measure, microplates were shaken for 400 sec with orbital shaking (87.6 rpm) and 200 sec with a linear shaking (135.6 rpm) to ensure proper yeast suspension and accurate measure.

The evolution of OD according to time was modeled by local polynomial regression fitting with the R-loess function setting the span parameter to 0.45. The growth ratio corresponds to the top OD rate in 1-hour windows in the conditions of interest normalized by those obtained on glucose media.

## Data availability

All sequencing data generated in this study have been deposited in the European Nucleotide Archive under the accession number PRJEB29566 for Illumina reads and PRJEB48853 for MinION reads.

## Acknowledgments

We are grateful to the IT and Bioinformatics core facility of the Institut de biologie moléculaire des plantes (IBMP-CNRS, Strasbourg, France) for providing computing and storage resources. This work was supported by the Agence Nationale de la Recherche (ANR-18-CE12-0013-02) and the European Research Council (ERC Consolidator Grant 772505). This work of the Interdisciplinary Thematic Institute IMCBio, as part of the ITI 2021-2028 program of the University of Strasbourg, CNRS, and Inserm, was supported by IdEx Unistra (ANR-10-IDEX-0002) and by SFRI-STRATUS project (ANR-20-SFRI-0012) and EUR IMCBio (ANR-17-EURE-0023) under the framework of the French Investments for the Future Program. J.-S.G. was supported by a grant from the French “Ministère de l’Enseignement Supérieur et de la Recherche.” J.S. is a Fellow of the University of Strasbourg Institute for Advanced Study (USIAS) and a member of the Institut Universitaire de France.

## Notes

### Competing Interest Statement

The authors have declared no competing interest.

