## Supplementary figures for "Contrasting genomic evolution between domesticated and wild *Kluyveromyces lactis* yeast populations"

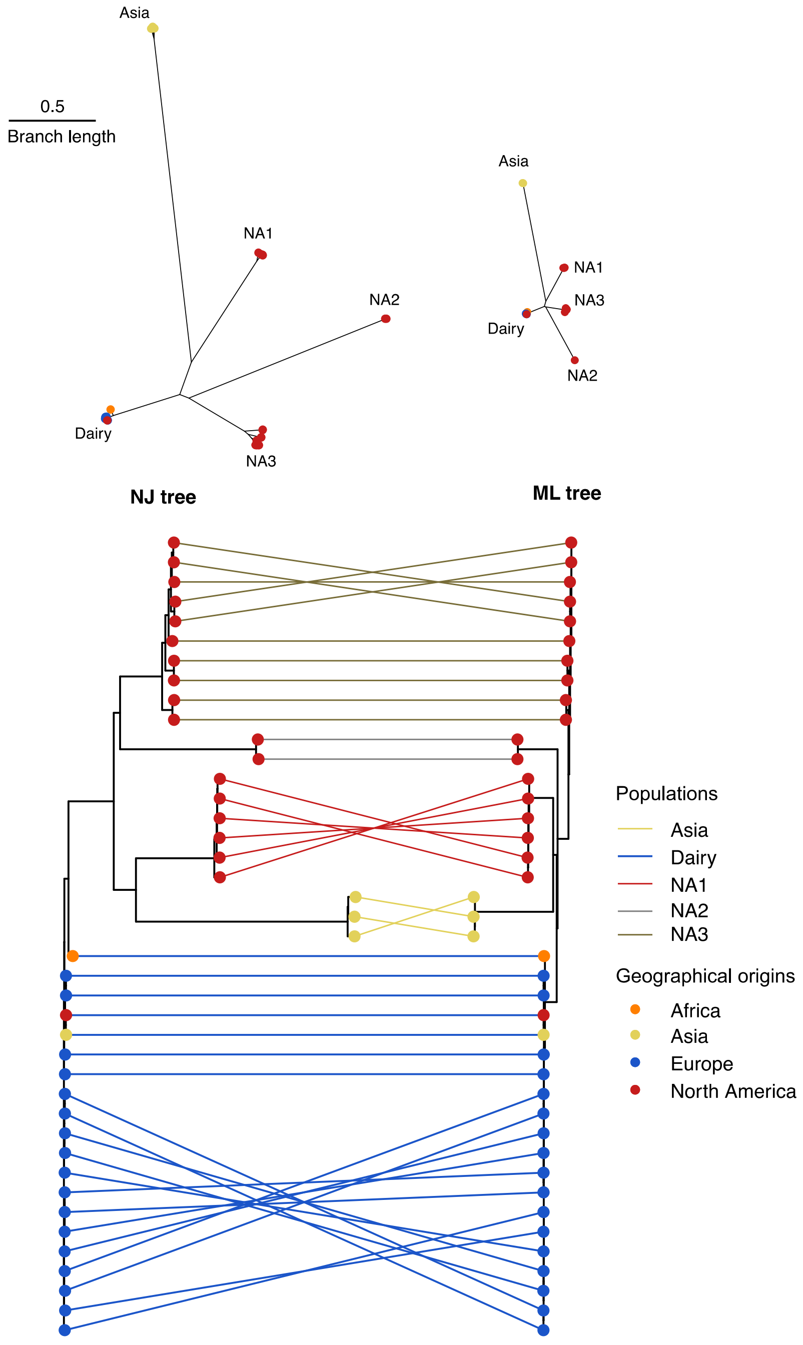


**Figure S1.** Comparison of the trees constructed with the whole SNP dataset based on a distance method (neighbor joining, NJ) or a maximum likelihood method (ML).
The tree comparison was obtained with R phytools library. The color of the nodes represents the geographical origins of the strains while the color of the link between the nodes is related to the population to which the strains belong.

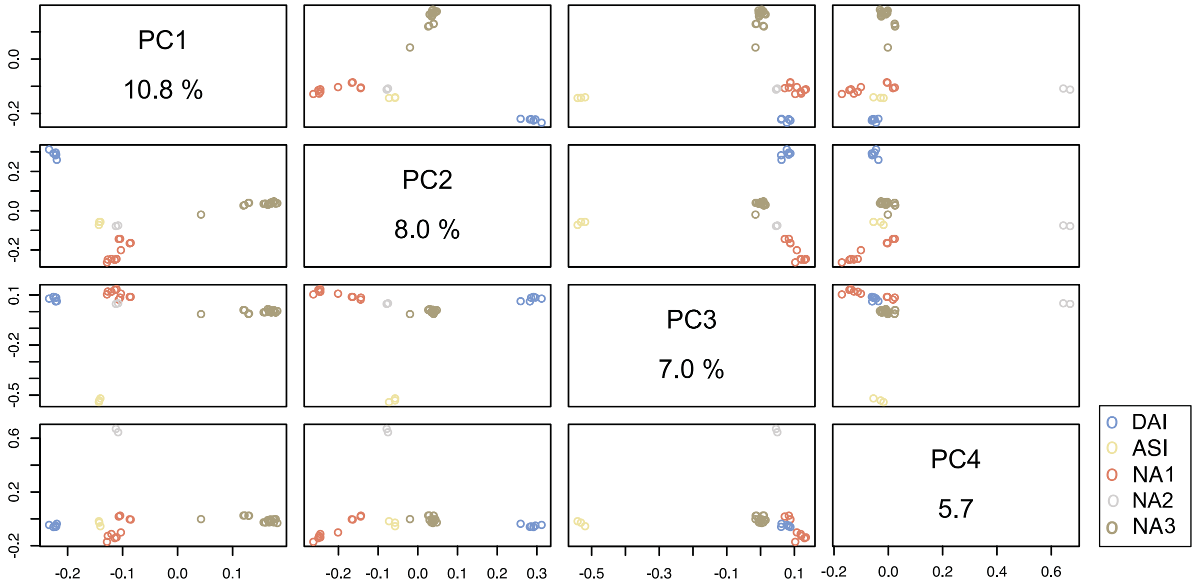


**Figure S2.** Principal-component analysis based on the whole genome SNP data of the 41 *Kluyveromyces lactis* sequenced strains.

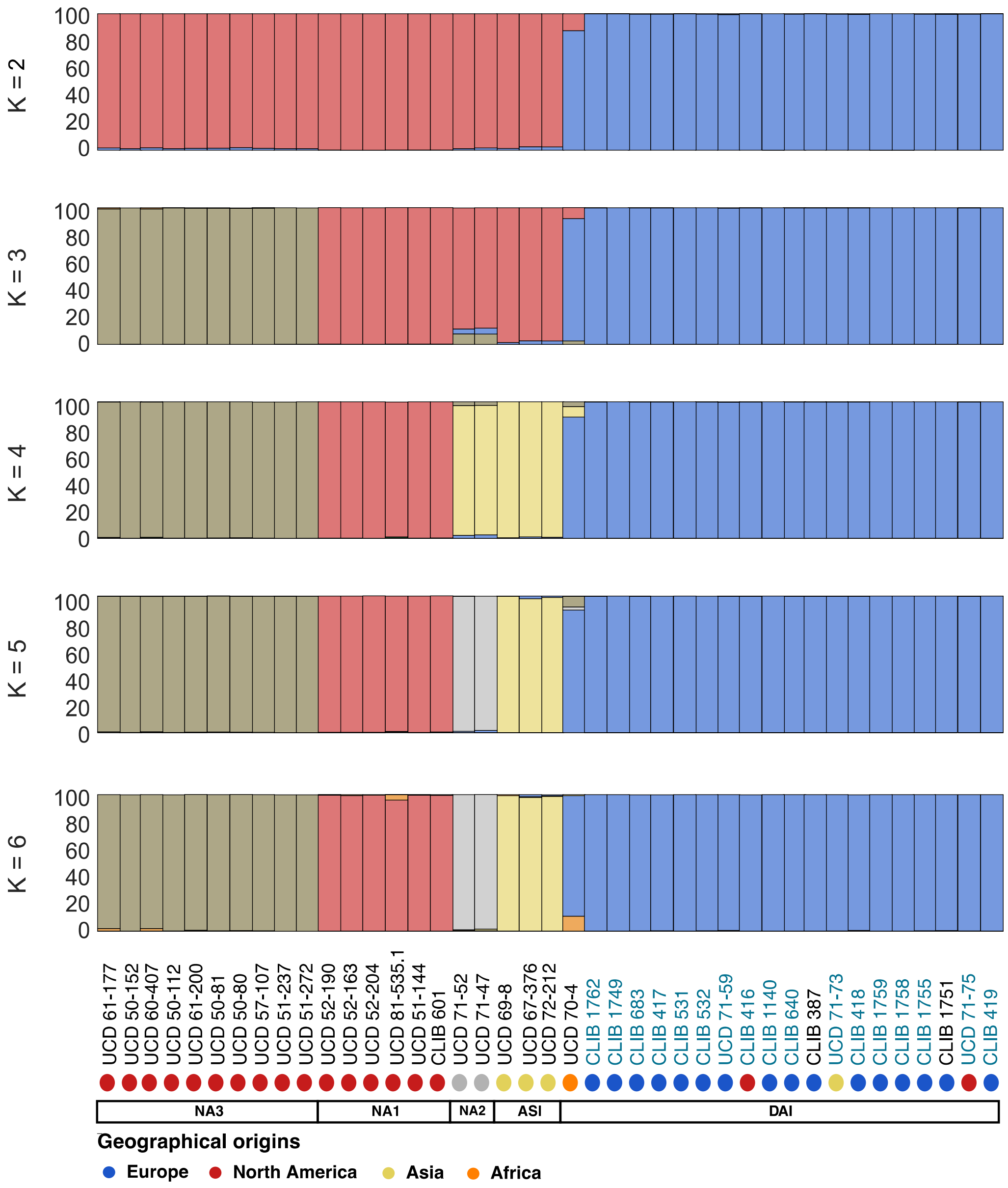


**Figure S3.** Structure of the *Kluyveromyces lactis* population. The number of populations (K) was predefined from 2 to 6. Each strain is represented by a single vertical bar, which is partitioned into K colored segments that represent the strain's estimated ancestry proportion in each of the K clusters.

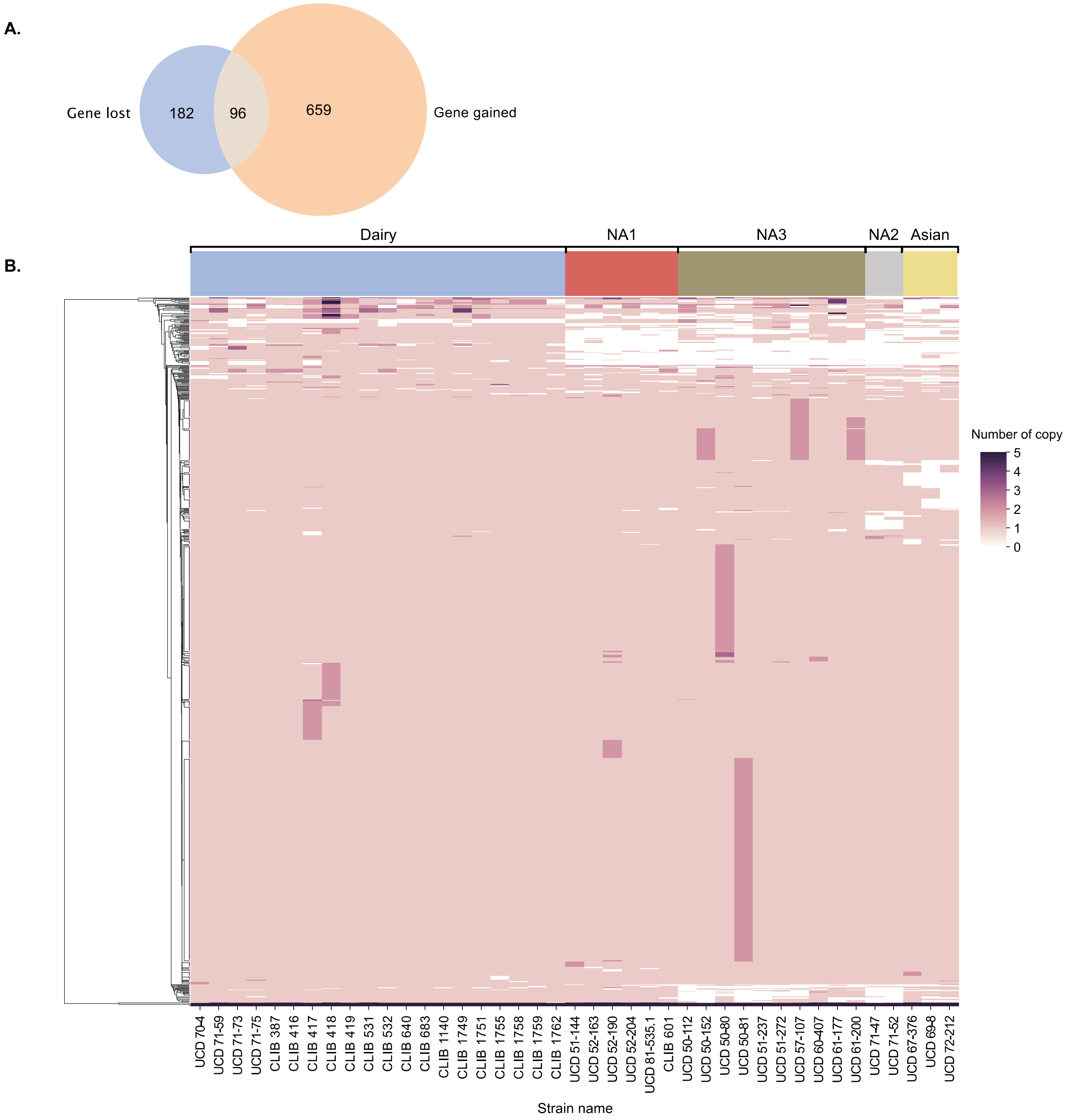


**Figure S4.** Copy number variant within the *Kluyveromyces lactis* species. The CNV detection was performed on the basis of the *Kluyveromyces lactis* var. *lactis* reference sequence. **A.** Venn diagram representing the number of unique genes that were detected as gained or lost within the species. **B.** Complete pattern of gene gain and loss in the 41 sequenced strains. The upper colors refer to the population to which the strains belong, as represented in Figure 1B.

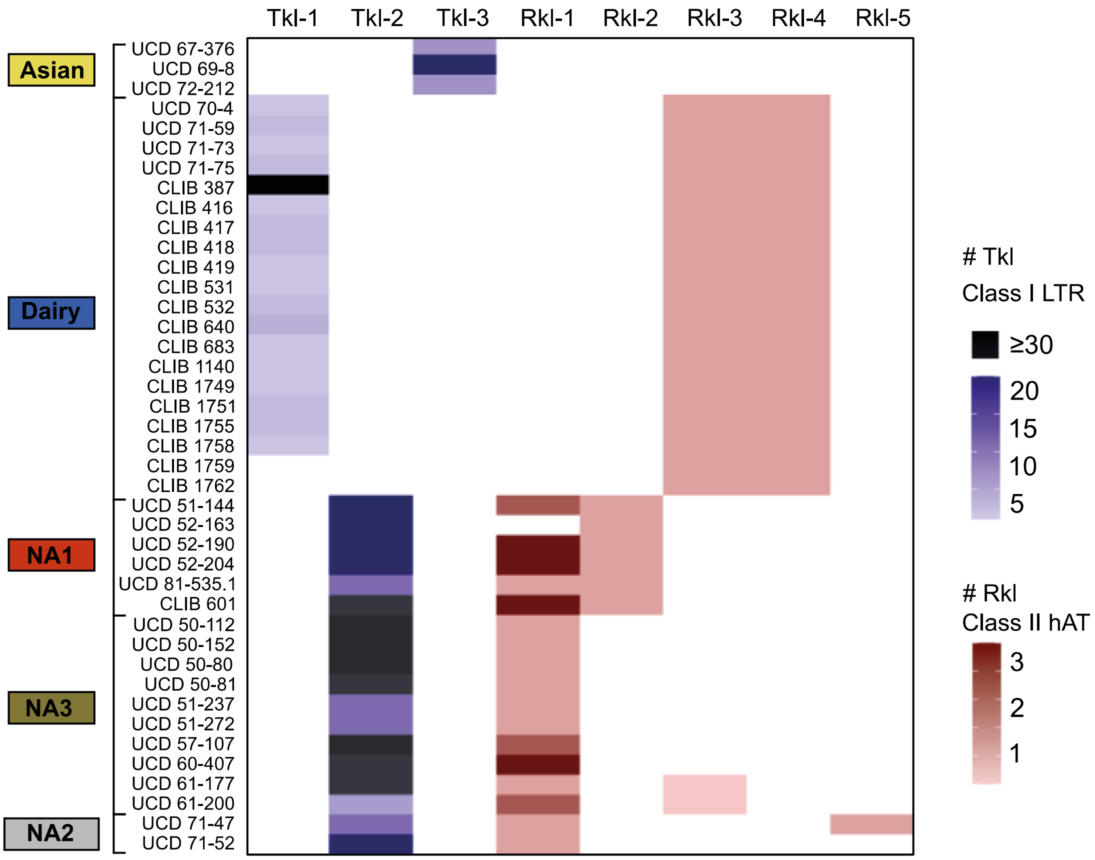


**Figure S5.** Transposable element content of the *Kluyveromyces lactis* species. The heatmap represents the number of copies of the transposable element variants for each of the 41 studied isolates. Strains are organized by population.

TkI = Ty *Kluyveromyces lactis*

RkI = Rover *Kluyveromyces lactis*

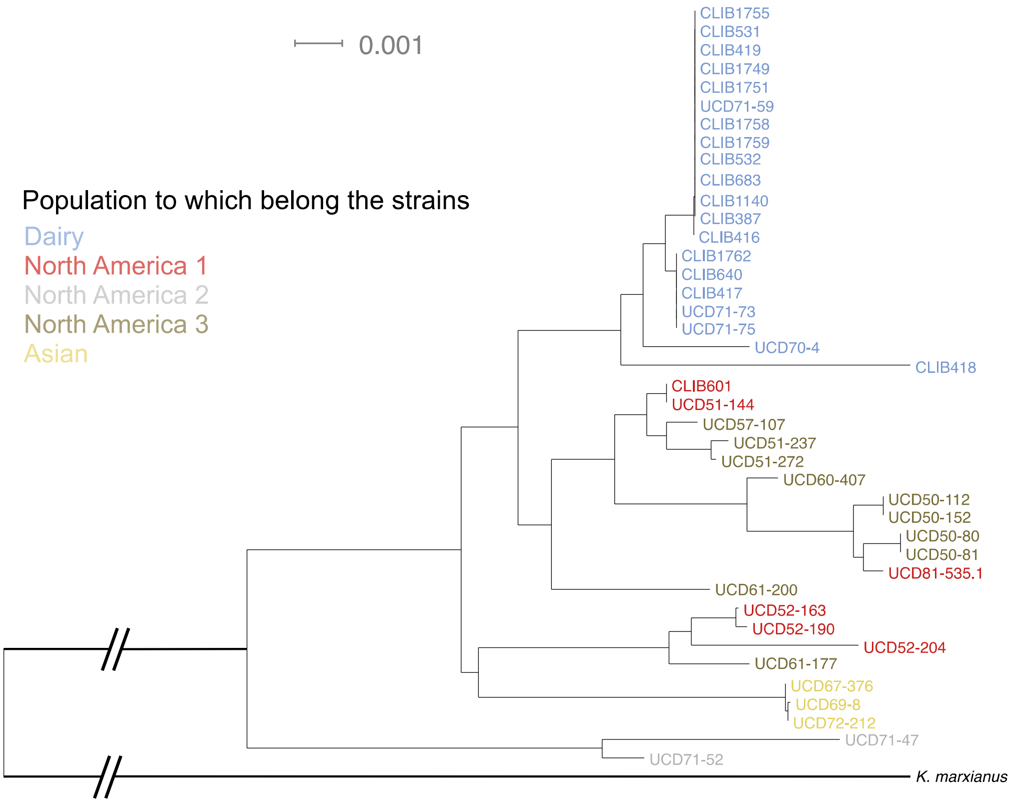


**Figure S6.** Neighbor-joining tree of mitochondrial sequences. This tree was constructed with BioNJ algorithm, on the basis of the concatenated coding sequences for *ATP6-COX2-COX3-CYTB*.
